# Metformin protects against insulin resistance induced by high uric acid in cardiomyocytes via AMPK signaling pathways in vitro and in vivo

**DOI:** 10.1101/2021.01.29.428905

**Authors:** Zhenyu Jiao, Yingqun Chen, Yang Xie, Yanbing Li, Zhi Li

**Author notes:** Corresponding author: Yanbing Li and Zhi Li. Zhenyu Jiao and Yingqun Chen contributed equally to this work.

## Abstract

High uric acid (HUA) is associated with insulin resistance and abnormal glucose metabolism in cardiomyocytes. Metformin is a recognized agonist of AMP-activated protein kinase (AMPK) and an antidiabetic drug widely used for type 2 diabetes. It can play a cardioprotective role in many pathways. We investigated whether metformin protects against HUA-induced insulin resistance and abnormal glucose metabolism in cardiomyocytes. We exposed primary cardiomyocytes to HUA, and cellular glucose uptake was quantified by measuring the uptake of 2-NBDG, a fluorescent glucose analog, after insulin excitation. Treatment with metformin (10 μmol/L) protected against HUA-inhibited glucose uptake induced by insulin in primary cardiomyocytes, as shown by fluorescence microscopy and flow cytometry analysis. HUA directly inhibited the phosphorylation of Akt and the translocation of glucose transporter type 4 (GLUT4) induced by insulin, which was blocked by metformin. Metformin promoted phosphorylation of AMPK, renewed HUA-inhibited glucose uptake induced by insulin and protected against insulin resistance in cardiomyocytes. As a result of these effects, in a mouse model of acute hyperuricemia, metformin improved insulin tolerance and glucose tolerance, accompanied by increased AMPK phosphorylation, Akt phosphorylation and translocation of GLUT4 in myocardial tissues. As expected, AICAR, another AMPK activator, had equivalent effects to metformin, demonstrating the important role of AMPK activation in protecting against insulin resistance induced by HUA in cardiomyocytes. Metformin protects against insulin resistance induced by HUA in cardiomyocytes and improves insulin tolerance and glucose tolerance in an acute hyperuricemic mouse model, along with the activation of AMPK. Consequently, metformin may be an important potential new treatment strategy for hyperuricemia-related cardiovascular disease.

## Introduction

Insulin resistance usually refers to a clinicopathological condition characterized by an impaired ability of insulin to stimulate glucose uptake and by glucose intolerance (Tan et al,. 2011). Moreover, insulin resistance is closely related to several cardiovascular diseases, such as myocardial infarction, hypertension and heart failure (Kanellakis et al., 2020; Wiebe et al., 2019; Uddin et al., 2019). Clinical follow-up studies have demonstrated that elevated serum uric acid is a strong independent risk factor for insulin resistance (Han et al., 2017), and ours and others’ previous studies have revealed that high uric acid (HUA) evokes insulin resistance in cardiomyocytes, adipocytes, muscle and liver (Zhi et al., 2016; Su et al., 2020; Zhu et al., 2014; Yuan et al., 2017).

Adenosine 5’-monophosphate-activated protein kinase (AMPK) is an essential metabolic regulator for energy sensing and has been well documented (Toyama et al., 2016); AMPK is expressed specifically in various tissues and cells, including cardiomyocytes (Hua et al., 2018), and plays a crucial role in regulating energy metabolism and energy homeostasis under stress conditions. AMPK can be activated by various conditions. Activation of AMPK phosphorylation regulates a series of metabolic steps by inducing the ATP generation system (such as glycolysis and fatty acid oxidation) and inhibiting the ATP consumption system (such as the synthesis of fatty acids). In addition, AMPK is very beneficial for improving insulin sensitivity in many cells and tissues. Our previous data demonstrated that AMPK is activated by HUA via oxidative stress in β cells (Zhang et al., 2013). In our previous study, HUA induced oxidative stress and played a critical role in the development of insulin resistance and its potential role in the pathogenesis of metabolic stress in cardiomyocytes (Zhi et al., 2016). Moreover, phosphorylated Akt-mediated insulin signal transduction and translocation of glucose transporter type 4 (GLUT4) were suppressed by HUA and were related to insulin resistance and impaired glucose uptake in cardiomyocytes (Zhi et al., 2016).

Metformin, well known as an AMPK activator and as an antidiabetic medicine, is widely used and has various pharmacologic effects, including promoting glucose uptake and utilization, enhancing insulin sensitivity and improving hyperlipidemia (Forslund et al., 2017). Major advances in the scientific understanding of metformin action have focused on the discovery that metformin leads to the activation of phospho-AMPK. This activation of phospho-AMPK appears to be more associated with changes in phosphocreatine levels and the AMP/ATP ratio. The AMPK system acts as an energy-sparing sensor, and therefore AMPK is a good mediator of metformin activity on heightened glucose uptake and the improved cellular status of energy that follows glucose uptake.

In our previous study, we found that HUA induced insulin resistance and suppressed glucose uptake in cardiomyocytes in vitro and in vivo (Zhi et al., 2016). In the present study, we assessed whether there are positive effects of metformin’s performance on glucose uptake inhibited by HUA in cardiomyocytes and whether treatment with metformin can either circumvent or reverse insulin resistance induced by HUA. At the cellular level, insulin resistance can be induced by prolonged incubation with HUA in cardiomyocytes. The impacts of HUA may be related to reductions in insulin signaling events at the level of phosphatidylinositol 3-kinase activation of Akt.

Previous studies have demonstrated that metformin can protect the heart by inhibiting apoptosis, autophagy, inflammation and other pathways via AMPK (Yuan et al., 2017; Hua et al., 2018; Forslund et al., 2017). However, whether metformin protects against insulin resistance induced by HUA in cardiomyocytes remains unknown. Therefore, we determined whether metformin activation of the AMPK-dependent pathway can protect against insulin resistance induced by HUA in cardiomyocytes.

## Methods & Materials

### Reagents

Uric acid (15 mg/dl, final concentration), metformin, insulin (100 nM, final concentration) and compound C (20 μM, final concentration) were purchased from Sigma (St. Louis, MO, USA). N-(7-Nitrobenz-2-oxa-1,3-diazol-4-yl)amino]-2-deoxy-d-glucose (2-NBDG, 100 μM, final concentration) and anti-IRS1 (Ser307) and anti-phospho-IRS1 (Ser307) antibodies were purchased from Invitrogen (Carlsbad, CA, USA). 5-Amino-4-imidazole-1-β-D-carboxamide ribofuranoside (AICAR, 500 μM, final concentration) was purchased from MedChemExpress (MCE LLC, USA). A rabbit anti-GAPDH antibody was purchased from Abcam (Abcam, USA). Anti-Akt (Ser473) and anti-phospho-Akt (Ser473) antibodies were purchased from Bioworld (St. Louis Park, USA). Anti-AMPK (Thr172) and anti-phospho-AMPK (Thr172) antibodies were purchased from Cell Signaling Technology (CST, Beverly, MA, USA). Cardiomyocytes were cultured in serum-free media for 24 hours and then incubated in the presence of 15 mg/dl HUA for 24 hours. Cardiomyocytes were pretreated with either metformin (0 to 20 μmol/L) or AICAR, an AMPK activator (500 μmol/L), for 60 minutes before the addition of HUA. Other cells were preincubated with compound C for 6 hours before the addition of either metformin or AICAR. Then, 2-NBDG uptake was analyzed in cardiomyocytes. were from Millipore (Billerica, MA).

### Cell culture

Cellular studies were carried out on primary cardiomyocytes isolated from C57BL/6 neonatal mice according to a previous protocol (Yue et al., 2012). Cardiomyocytes were incubated with Dulbecco’s modified Eagle medium (DMEM) or low-glucose minimum essential medium eagle (MEM) containing L-glutamine (2 mM), streptomycin (100 mg/ml), penicillin G (100 U/ml) and 10% fetal bovine serum. For all experiments, the cardiomyocytes were seeded into 6-well plates at a density of 2.0×10^5^ cells/ml. Primary cardiomyocytes were incubated in a 37°C incubator with 95% air and 5% CO_2_.

### Uptake of 2-NBDG glucose was measured by fluorescence microscopy and flow cytometry

A fluorescent glucose analog, 2-NBDG, glucose uptake, was measured by fluorescence microscopy and flow cytometry in cardiomyocytes to assess glucose uptake.

Briefly, for fluorescence microscopy, the cells were incubated in fetal bovine serum-free medium for 24 hours. The culture medium was replaced by Krebs-Ringer-Bicarbonate buffer containing insulin (100 nM) and 2-NBDG (100 μM) at 37°C for 30 minutes and analyzed at excitation and emission wavelengths of 488 and 525 nm, respectively.

For flow cytometry, the cells were fed low-glucose DMEM without fetal bovine serum for 24 hours. The culture medium was then replaced with Krebs-Ringer-Bicarbonate (KRB) buffer containing 2-NBDG (100 μM) and insulin (100 nM) for 30 minutes at 37°C in a CO_2_ incubator. For flow cytometry-based 2-NBDG glucose uptake assays, after treatment, 2-NBDG was washed out of the culture medium, and uptake of 2-NBDG was assessed by flow cytometry with a fluorometer at an excitation wavelength of 488 nm and an emission wavelength of 525 nm. For each sample, 20,000 cells in each well were obtained in the FSC x SSC plots.

### Animal experiments

The protocol was unanimously affirmed by the Shantou University Animal Experiments Ethics Committee. All methods were performed in accordance with the relevant guidelines. C57BL/6J male mice (8 weeks old) were obtained from Vital River Laboratories Animal, Ltd. (Beijing, China) in this study. All animals were housed in the Laboratory Animal Center of Shantou University Medical College. The animals were housed in cages under a 12-hour light-dark cycle and fed water and a normal food diet. Cardiac myocytes and ventricle cardiac tissue were obtained as previously reported (Wang et al., 2013).

### Experimental acute hyperuricemia mouse model establishment and metformin treatment

Ten-week-old male mice were randomly divided into 3 groups (n=6 each) for treatment: the control, HU (hyperuricemia) and HU-Met (metformin) groups. For the HU group, following an 18-hour overnight fast, the mice were injected intraperitoneally (i.p.) with 300 mg/kg potassium oxonate and intragastrically with 500 mg/kg hypoxanthine to create the model of acute hyperuricemia for 1-2 hours. The mice in the HU-Met group were administered 250 mg/kg/day metformin in drinking water for 14 consecutive days (Han et al., 2017). The quantity of medicine was based on body weight levels before each dose. The level of serum uric acid was determined by the phosphotungstic acid method at different times (Li et al., 2011). Insulin tolerance tests (ITTs) and glucose tolerance tests (GTTs) were performed as described previously (Zhu et al., 2014). For the control group and HU group, the mice were sacrificed by CO_2_ inhalation 10 minutes after injection with 2 U/kg insulin. Cardiac muscle tissue was excised and stored in liquid nitrogen immediately.

### Western blot

Cardiomyocytes were washed twice with cold PBS, harvested, lysed in ice-cold lysis buffer (pH=7.4, 100 mM KCl, 10 mM HEPES, 1.5 mM MgCl), sonicated and homogenized in RIRP buffer. After centrifugation, a Pierce BCA protein assay kit was used to determine the protein concentration. Equal concentrations of soluble lysate protein (50 mg) were added to a 12% SDS-PAGE Bis-Tris gel, and the protein was transferred to immunoblot polyvinylidene fluoride (PVDF) membranes (Bio-Rad, USA). Membranes were incubated and blocked overnight at 4°C with antibodies against p-AMPK/AMPK, p-Akt/Akt and GAPDH. Then, the membranes were incubated with anti-rabbit horseradish peroxidase-conjugated secondary antibodies (1:10,000 dilution). The immunoreactive bands were analyzed with an enhanced chemiluminescence substrate kit (Biological Industries, BI, Israel).

### Statistical analysis

Data are presented as the means ± SD. Significant differences between the means were analyzed by a Student’s t-test. Significant differences were evaluated using one-way analyses of variance (ANOVA), and P values less than 0.05 were considered to be statistically significant.

## Results

### Metformin stimulated the phosphorylation of AMPK and attenuated insulin resistance induced by HUA in cardiomyocytes

We first wanted to determine the most suitable exposure period and concentration of metformin for the activation of phospho-AMPK in cardiomyocytes. Treatment with metformin (using the most suitable concentration and exposure period of 10 μM for 60 minutes) stimulated AMPK phosphorylation in cardiomyocytes in a dose- and time-dependent manner (Figure 1A and 1B). Moreover, 2-NBDG glucose uptake was decreased significantly in the presence of HUA (15 mg/dl) for 24 hours in cardiomyocytes, as shown by the flow cytometry assay. These results demonstrated that HUA inhibited 2-NBDG glucose uptake induced by insulin and caused insulin resistance in primary cardiomyocytes, but this change was attenuated in a dose- and time-dependent manner by treatment with metformin (Figure 1C).

**Fig 1.**
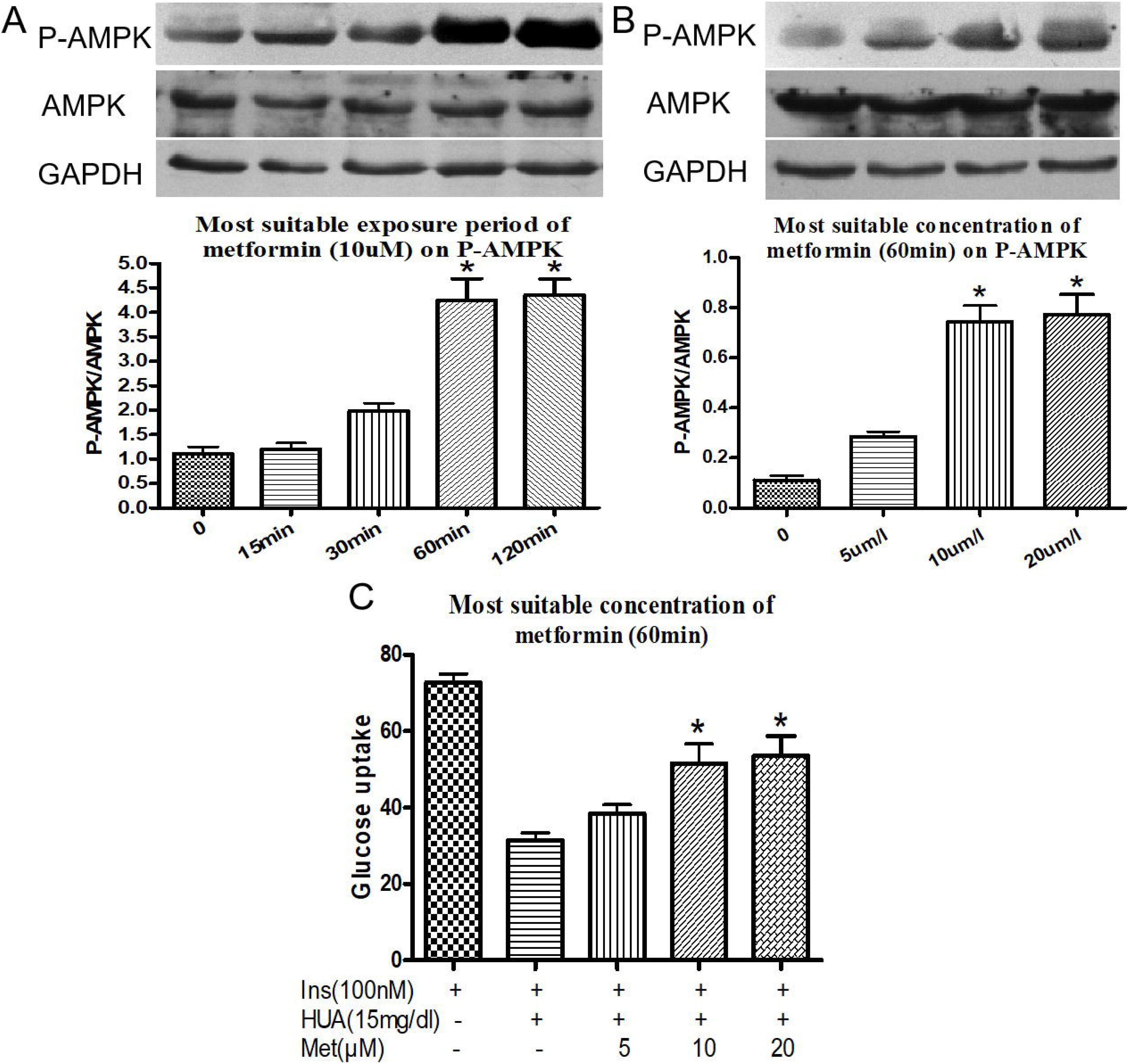
Effect of metformin on AMPK phosphorylation (A-B) and HUA-inhibited glucose uptake induced by insulin in primary cardiomyocytes (C). **(A-B)** Western blot analysis of phosphorylated and total AMPK levels. **(C)** Pretreated with metformin (10 μmol/L) for 60 minutes protected agains HUA (15mg/dl for 24 hours) inhibited 2-NBDG uptake induced by insulin in primary cardiomyocytes, as shown by flow cytometry analysis. **A,** *P<0.05 vs. 0, 15 and 30 min. **B,** *P<0.05 vs. 0 and 5uM. **C,**\*P<0.05 vs. Ins+HUA. Data are mean±SD from 4 separate experiments. Ins: insuln. HUA: high uric acid. Met: metformin. 2-NBDG: 2-[N-(7-Nitrobenz-2-oxa-1,3-diazol-4-yl)amino]-2-deoxy-d-glucose.

### Metformin attenuated insulin resistance induced by HUA via AMPK signaling pathways in cardiomyocytes

To investigate whether metformin attenuates HUA-induced insulin resistance via AMPK signaling pathways in cardiomyocytes, cardiomyocytes were pretreated with metformin (10 μM) for 60 minutes and/or compound-C (20 μM), a specific AMPK inhibitor, for 6 hours before exposure to HUA. Under these conditions, the effect of metformin on 2-NBDG glucose uptake was blunted by cotreatment with compound C (Figure 2A-2C).

**Fig 2.**
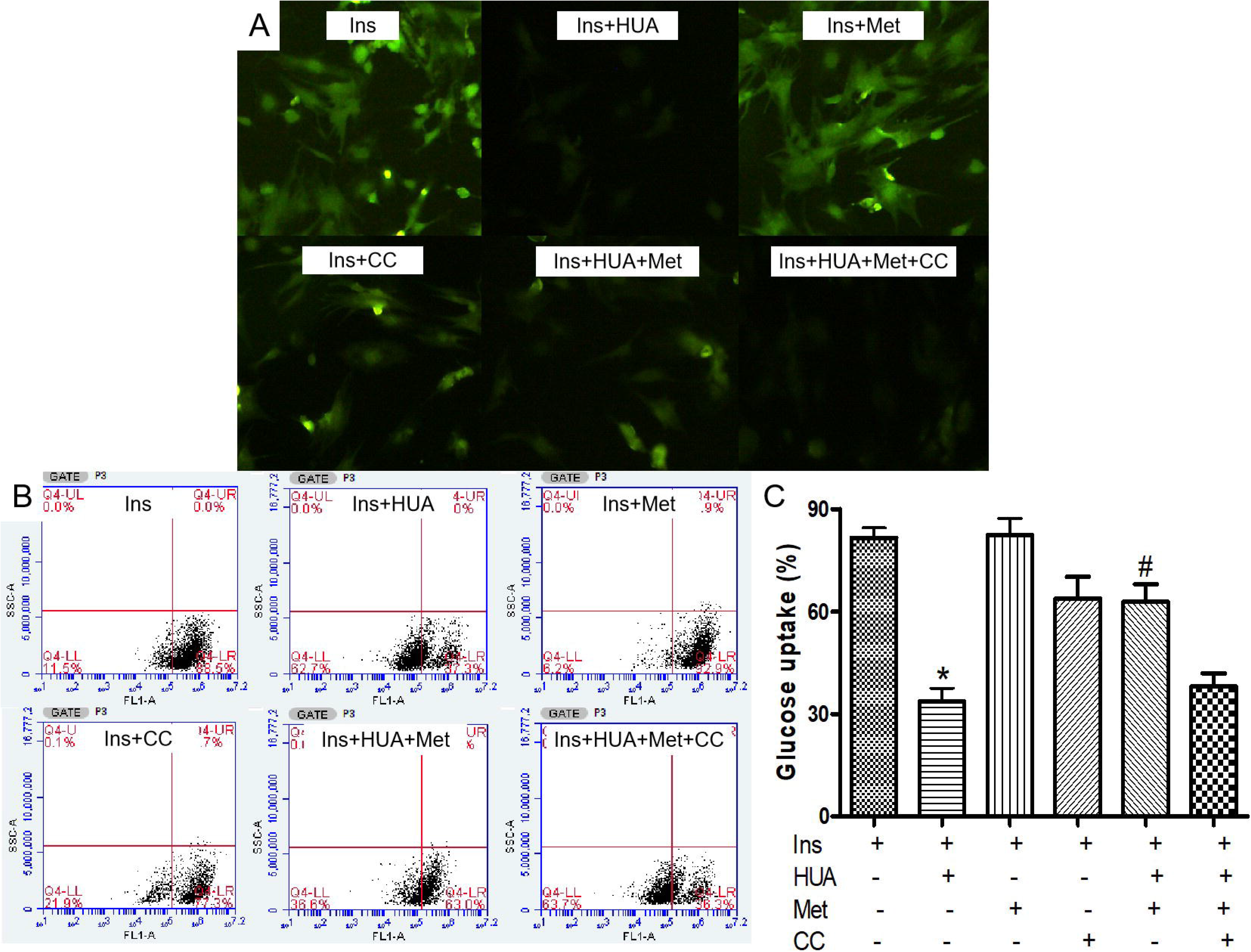
Metformin protected agains HUA inhibited 2-NBDG uptake induced by insulin via AMPK signal pathways in primary Cardiomyocytes. **(A)** 2-NBDG uptake assay detected by fluorescence microscopy. **(B-C)** 2-NBDG uptake assay detected by flow cytometry. *P<0.05 vs. Ins and Ins+HUA+Met, #P<0.01 vs. Ins+HUA+Met+CC. Data are mean±SD from 4 separate experiments. CC: compound C, an AMPK inhibitor.

To confirm the role of AMPK signaling pathways in metformin attenuation of HUA-induced insulin resistance in cardiomyocytes, cells were pretreated with another AMPK activator, AICAR (500 μM), for 60 minutes before exposure to HUA. We found that AICAR had an effect similar to metformin on glucose (2-NBDG) uptake after exposure to HUA (Figure 3A-3C). These findings demonstrated that activation of AMPK signaling pathways protected against insulin resistance induced by HUA in cardiomyocytes.

**Fig 3.**
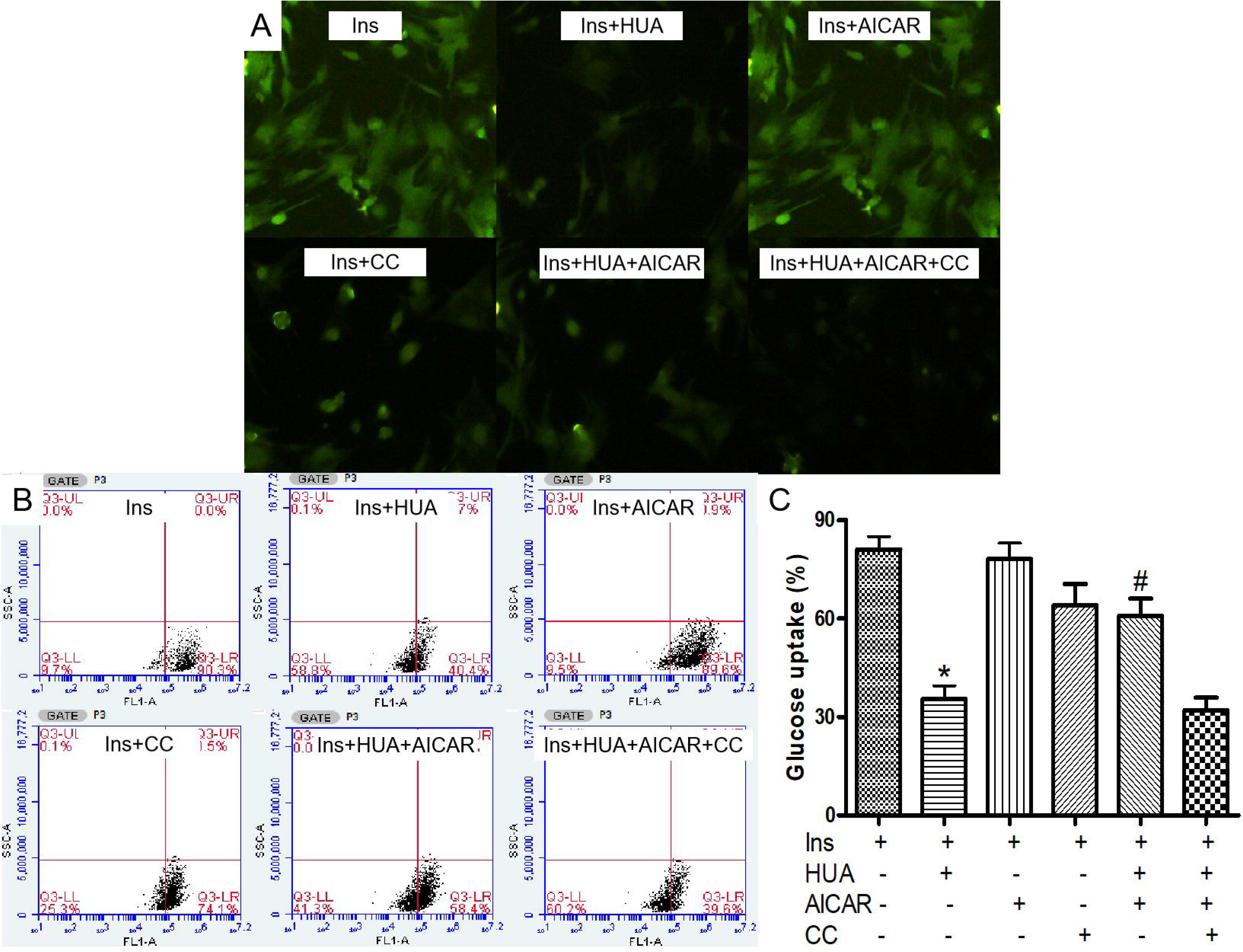
5-amino-4-imidazole-1-β-D-carboxamide ribofuranoside (AICAR, another AMPK activator) protected agains HUA inhibited 2-NBDG uptake induced by insulin via AMPK signal pathways in primary Cardiomyocytes. **(A)** 2-NBDG uptake assay detected by fluorescence microscopy. **(B-C)** 2-NBDG uptake assay detected by flow cytometry. *P<0.05 vs. Ins and Ins+HUA+AICAR, #P<0.01 vs. Ins+HUA+AICAR+CC. Data are mean±SD from 4 separate experiments.

### Metformin increased phospho-AMPK, phospho-Akt and translocation of GLUT4 but had no effect on phospho-IRS1 expression in cardiomyocytes

AMPK is a well-known master energy sensing regulator. Furthermore, the Akt family of Thr/Ser protein kinases is of critical importance with regard to cardiac metabolism and growth in insulin signal transduction. Consequently, to further inspect the molecular mechanism of the AMPK and Akt signaling pathways, we first examined whether metformin activated AMPK in cardiomyocytes. In cardiomyocytes, metformin enhanced the phosphorylation of AMPK, and this phosphorylation was blocked by compound C, a specific AMPK inhibitor (Figure 4A).

**Fig 4.**
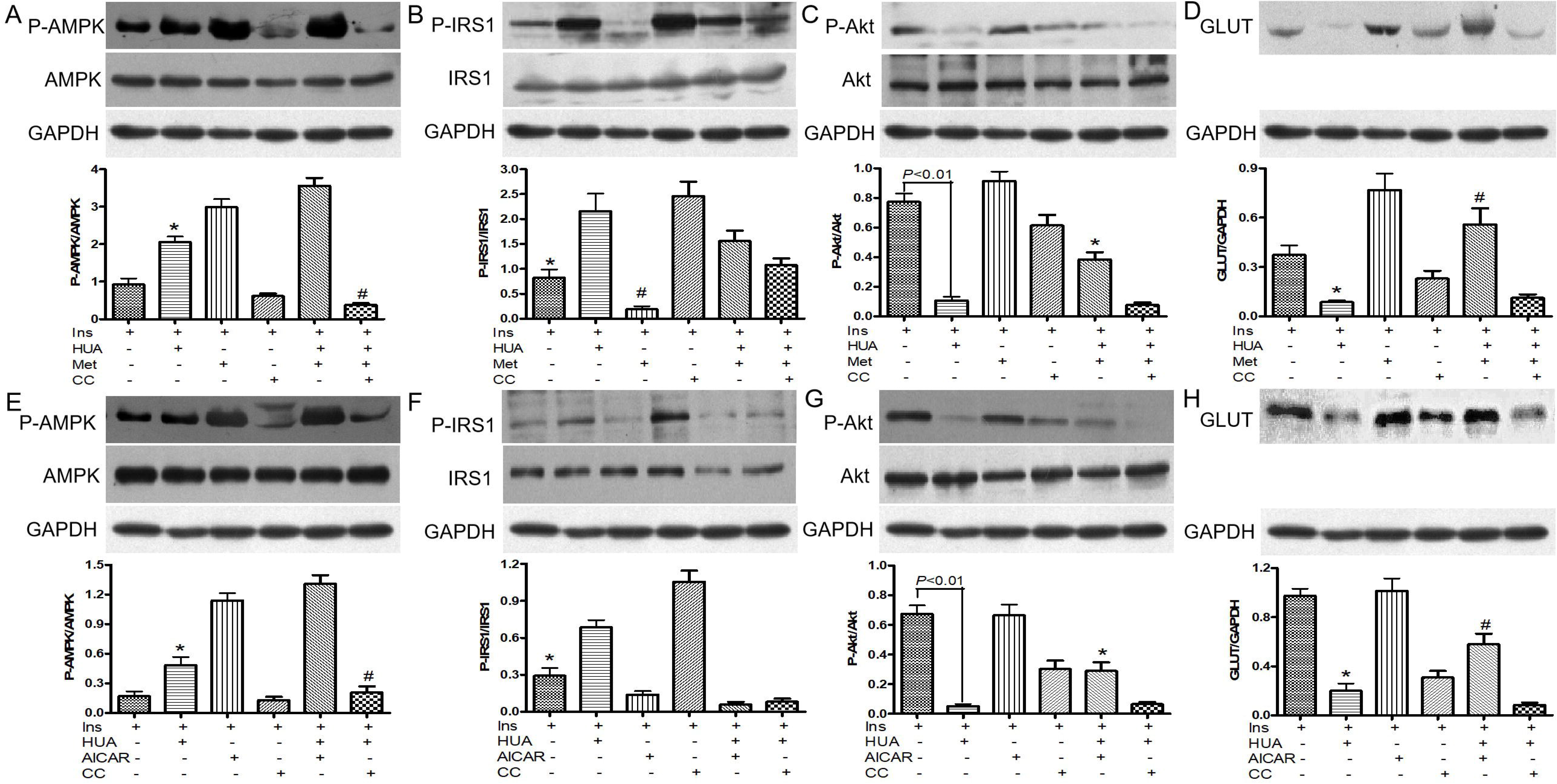
(A-D) Effect of merformin on HUA-induced/inhibited of phospho-AMPK (A), phospho-IRS1 (B), phospho-Akt (C) and GLUT (D) in primary Cardiomyocytes. **A,** *P<0.05 vs. Ins and Ins+HUA+Met; #P<0.01 vs. Ins+HUA+Met. **B,** *P<0.05 vs. Ins+HUA and Ins+CC; #P<0.01 vs. Ins+HUA+Met. **C,** *P<0.05 vs. Ins+HUA, Ins+Met and Ins+HUA+Met+CC. **D,** *P<0.05 vs. Ins and Ins+HUA+Met; #P<0.01 vs. Ins+HUA. **(E-H) Effect of AICAR on HUA-induced/inhibited of phospho-AMPK (E), phospho-IRS1 (F), phospho-Akt (G) and GLUT (H) in primary Cardiomyocytes. E,** *P<0.05 vs. Ins and Ins+HUA+Met; #P<0.01 vs. Ins+HUA+Met. **F,** *P<0.05 vs. Ins+HUA and Ins+CC. **G,** *P<0.05 vs. Ins+HUA, Ins+Met and Ins+HUA+Met+CC. **H,** *P<0.05 vs. Ins and Ins+HUA+Met; #P<0.01 vs. Ins+HUA.

To test whether metformin also activated the expression of phospho-Akt and translocation of GLUT4 in cardiomyocytes, we determined phospho-Akt expression and translocation of GLUT4 by performing western blot analyses in cardiomyocytes. We demonstrated that the insulin-induced phosphorylation level of Akt and translocation of GLUT4 were significantly inhibited by HUA (Figure 4B-4C) but had no effect on phospho-IRS1 (Figure 4D) expression in cardiomyocytes. Furthermore, we investigated the role of metformin in HUA-mediated inhibition of Akt activation and GLUT4 translocation. Metformin did not influence the insulin-induced activation of Akt kinase phosphorylation or translocation of GLUT4 (Figure 4B-4C). However, insulin-induced activation of Akt kinase phosphorylation and translocation of GLUT4 were depressed by HUA, which was attenuated significantly by metformin (Figure 4B-4C). Nevertheless, the effect of metformin on Akt phosphorylation and translocation of GLUT4 were depressed by HUA, which was significantly offset by compound C, an AMPK inhibitor. As expected, we found that AICAR had an effect similar to metformin on phospho-AMPK, phospho-Akt and translocation of GLUT4 after exposure to HUA (Figure 4A-4C). These results demonstrated a profound effect of metformin on downstream targets of the insulin signaling pathway via AMPK.

### Effect of metformin in a hyperuricemia mice model

#### Effect of metformin on insulin resistance induced by hyperuricemia in a mouse model

As expected, the levels of serum uric acid in the acute hyperuricemia mouse model were significantly higher than those before hyperuricemia induction (115.69±18.32 vs 39.71± 4.43 mg/l), which was consistent with primary hyperuricemia patients (Nakagawa et al., 2006). Our acute hyperuricemia mouse model also indicated an impaired glucose tolerance test (Figure 5A) and insulin tolerance test (Figure 5B) at 15 and 30 minutes after insulin or glucose injection with insulin resistance. Treatment with metformin significantly decreased the level of serum glucose in the glucose tolerance test and insulin tolerance test compared with the hyperuricemia mouse model (Figure 5A and 5B). Thus, these findings demonstrate that metformin protects against hyperuricemia-induced insulin resistance in a mouse model.

**Fig 5.**
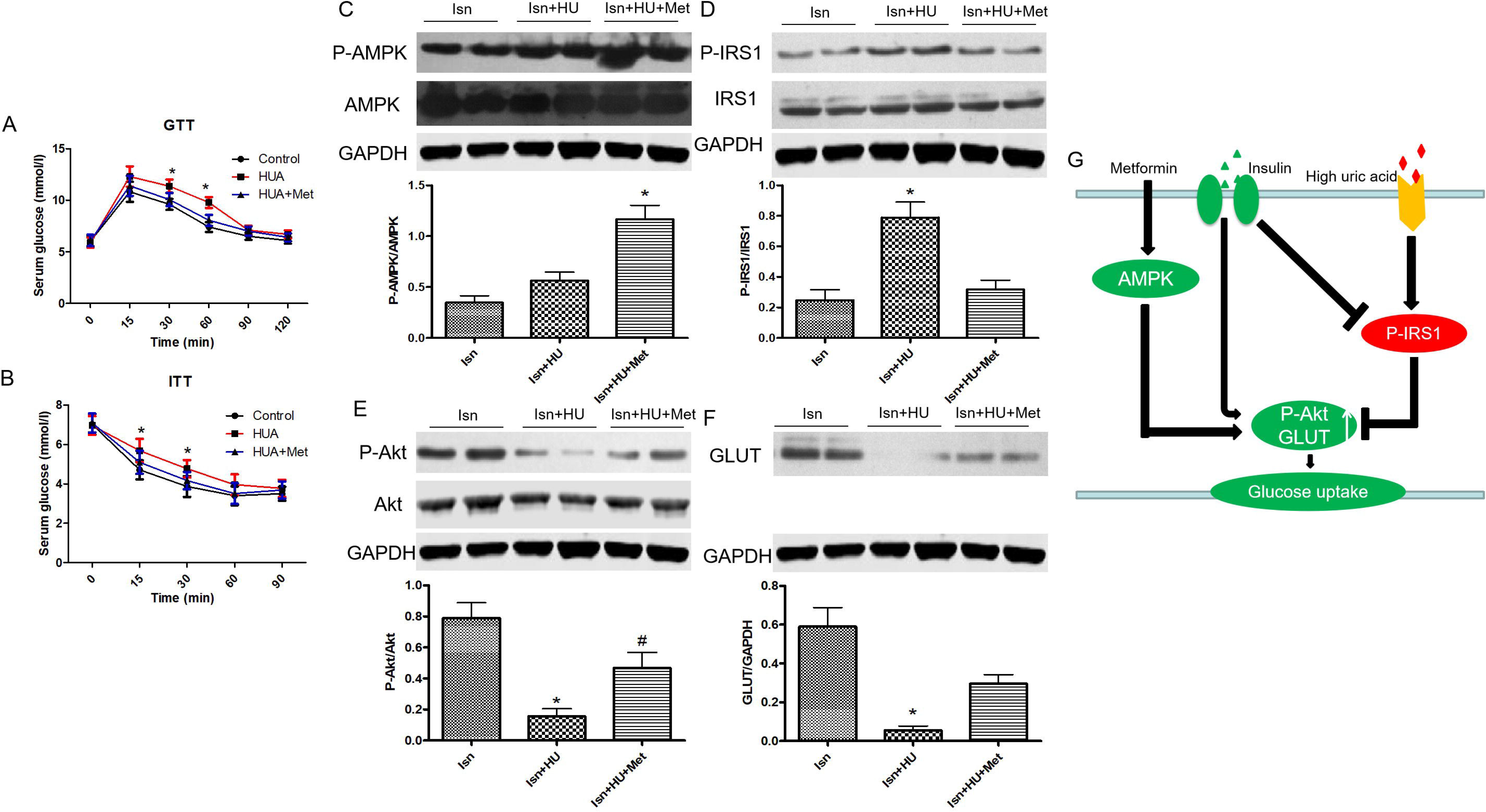
(A-B) Glucose tolerance test (GTT) (A) and insulin tolerance test (ITT) (B) in an acute hyperuricemic mice model. *P < 0.05 vs. Con and HU+Met. **(C-F) Western blot analysis of phospho-AMPK (C), phospho-IRS1 (D), phospho-Akt (E) and GLUT (F) level in cardiac tissues. C,** *P < 0.05 vs. Ins+HU. **D,**\*P < 0.05 vs. Ins and Ins+HU+Met. **E,** *P < 0.05 vs. Ins and Ins+HU+Met. **F,** *P < 0.05 vs. Ins and Ins+HU+Met. Data are mean ± SD from 4 separate experiments. (**G) Schematic representation of how metformin ameliorates HUA-induced insulin resistance in cardiomyocytes**. After treatment with HUA, which activates phosphorylation of IRS-1 (Ser307). This activity impairs AKT (Ser473) phosphorylation, for insulin resistance. After treatment with metformin, the phosphorylation of AMPK is increased. At the same time, this activity reverses the inhibition of HUA-induced AKT phosphorylation, which ameliorates HUA-induced insulin resistance. HU: hyperuricemia. Met: metformin.

#### Effect of metformin on AMPK, IRS1 phosphorylation, Akt phosphorylation and translocation of GLUT4 in cardiac tissues from the acute hyperuricemic mouse model

To examine the molecular mechanisms by which metformin protects against hyperuricemia-induced insulin resistance in cardiomyocytes in an acute hyperuricemic mouse model, we investigated the effect of metformin on AMPK, IRS1 phosphorylation, Akt phosphorylation and GLUT4 translocation in cardiac tissues from the acute hyperuricemic mouse model. The hyperuricemic mice were injected with 2 U/kg insulin, and after 10 minutes, the cardiac tissues were isolated and obtained. Metformin increased the phosphorylation level of AMPK in mouse cardiac tissues (Figure 5C). The western blot analysis results demonstrated a significant reduction in the level of phospho-AKT and translocation of GLUT4 in the hyperuricemic mouse model, while metformin reversed these changes (Figure 5E-5F) but had no effect on phospho-IRS1 (Figure 5D). These results demonstrate that metformin protects against hyperuricemia-induced insulin resistance in cardiomyocytes via AMPK signaling pathways in vivo.

## Discussion

To the best of our knowledge, this is the first study to clearly illustrate that metformin, which is used worldwide as an antidiabetic medicine, protects against insulin resistance in cardiomyocytes induced by HUA and increases AMPK activation. Of course, we and others have previously shown that HUA induces insulin resistance in various cells, including cardiomyocytes, skeletal muscle cells and HepG2 cells (Zhi et al., 2016; Su et al., 2020; Zhu et al., 2014; Yuan et al., 2017). Nevertheless, it has been unclear whether AMPK can regulate insulin resistance induced by HUA in cardiomyocytes in vitro and in vivo. Therefore, we used neonatal mouse primary cardiomyocytes and an acute hyperuricemia mouse model, which is regarded as similar to humans with primary hyperuricemia and can be utilized in translational research for human hyperuricemia.

Hyperuricemia is strongly associated with cardiovascular risk and a poor outcome in a variety of cardiovascular disease states, such as coronary artery disease, hypertension and heart failure (Tai et al., 2020; Huang et al., 2020; Krishnan et al., 2006; Anker et al., 2003; Doehner et al., 2005). However, the underlying mechanisms to explain this association have been nonexistent. In our previous study (Zhi et al., 2016), we explored the impact of HUA on glucose uptake and insulin resistance in primary cardiomyocytes and found that HUA induces oxidative stress, which plays a crucial role in the progression of insulin resistance in cardiomyocytes. Moreover, recent studies have confirmed insulin resistance as a powerful independent predictor of mortality and morbidity in patients with heart failure (Szabo et al., 2011; Yang et al., 2006). These studies demonstrated that therapeutically targeting impaired insulin sensitivity may potentially be favorable for patients with chronic heart failure. To determine whether metformin protects against insulin resistance induced by HUA in cardiomyocytes, we exposed primary cardiomyocytes to HUA, pretreated them with metformin and then quantified the uptake of glucose with 2-NBDG, a fluorescent glucose analog, after insulin stimulation. We found that treatment with metformin may protect cardiomyocytes from HUA and inhibit insulin-induced glucose uptake in cardiomyocytes. These results suggested that HUA resulted in insulin resistance in cardiomyocytes, but this change was weakened by pretreatment with metformin.

Metformin is well known to activate AMPK, which is expressed in a variety of tissues and cells, including cardiomyocytes, and plays a pivotal role in the regulation of cellular energy metabolism under stress conditions (Sasaki et al., 2009). Previous studies have shown that metformin reduces long-term and high insulin-induced insulin resistance in cardiomyocytes (Vlavcheski et al., 2020), and AMPK has been shown to protect against insulin resistance in skeletal muscle cells through restoration of GLUT4 translocation (Zhang et al., 2018). Consistent with the findings of these previous studies, we found that metformin could improve HUA-induced insulin resistance in primary cardiomyocytes. As expected, this change was blocked by compound C, an AMPK inhibitor, indicating that AMPK activation was responsible for the suppression of insulin resistance in cardiomyocytes. In addition, using an acute hyperuricemic mouse model, our study indicated that metformin improved the progression of insulin resistance induced by HUA, as shown by the glucose tolerance test (Figure 5A) and insulin tolerance test (Figure 5B).

Interestingly, another activator of AMPK, AICAR, had almost the same effects as metformin, suggesting that activation of AMPK phosphorylation promoted the observed protective effect of insulin resistance in cardiomyocytes. In fact, AICAR has been demonstrated to protect against myocardial ischemia-reperfusion injury after myocardial infarction in animals and humans (Mangano., 1997; Beker et al., 2019). However, what procedures following AMPK pathway activation are involved in cardioprotection?

The first possibility is the improvement of the insulin-activated Akt pathway, which is inhibited by HUA. A previous study demonstrated that phosphorylation of Akt regulates cell survival, growth and metabolism. Additionally, cardiomyocyte metabolism and growth are coordinated by the integration of intracellular and extracellular signals (Russell et al., 1999). Furthermore, activation of Akt phosphorylation, a key protein kinase of insulin-induced glucose uptake, modulates glucose uptake stimulated by insulin in cardiomyocytes (Zhi et al., 2016). In the present study, we found that HUA could strongly inhibit Akt phosphorylation, translocation of GLUT4 and 2-NBDG glucose uptake induced by insulin in primary cardiomyocytes. Our findings suggest that activators of AMPK, such as metformin and AICAR, could prevent HUA-inhibited Akt phosphorylation, GLUT4 translocation and HUA-inhibited 2-NBDG glucose uptake in cardiomyocytes. Moreover, an acute hyperuricemia mouse model demonstrated inhibited Akt phosphorylation with insulin resistance and glucose intolerance.

Another possibility is the metabolic effects of AMPK phosphorylation activation. Metformin has been demonstrated to activate the phosphorylation of AMPK in cardiomyocytes and mouse cardiac tissues. In fact, a recent study suggested that short-term treatment with metformin protects against myocardial ischemia-reperfusion injury via the AMPK signaling pathway after myocardial infarction in mice (Hua et al., 2018). Therefore, AMPK has been considered to have various cardioprotective effects in animals and humans. Improvement in AMPK production by metformin may have mitigated the progression of insulin resistance induced by HUA. Both AICAR and metformin are reported to enhance glucose uptake in skeletal muscle cells and heart muscle cells (Yuan et al., 2017; Russell et al., 1999). Consistent with these reports, in the HUA plus insulin group, we found that pretreatment with metformin and AICAR nearly reverted glucose uptake to the level of the control group in cardiomyocytes. Therefore, the possibility exists that the AMPK-induced uptake of glucose triggers improvement in insulin resistance, which is induced by HUA, followed by the restitution of the metabolic switch.

The decreased Akt phosphorylation and translocation of GLUT4 in the HUA group were reversed in both the HUA plus AICAR group or the HUA plus metformin group, demonstrating that the activation of AMPK protects the insulin signaling pathway of Akt against inhibition by HUA in both the HUA plus AICAR group or the HUA plus metformin group.

## Conclusions

Our findings illustrated that metformin protects against the progression of insulin resistance induced by HUA in cardiomyocytes and activates phospho-AMPK. These findings show an interaction between activation of AMPK induced by metformin and insulin resistance induced by HUA that could be important to our understanding of the potentiating effects of metformin on patients with insulin resistance and hyperuricemia in clinical trials. Therefore, metformin may provide a new treatment strategy for hyperuricemia-related cardiovascular disease.

## Declarations

### Acknowledgments and funding support

This work was supported by grants from the Guangdong Basic and Applied Basic Research Foundation of China (2018A030307056), the Guangdong Medical Science and Technology Research Fund Project of China (A2020159) and the Shantou Science and Technology Plan Project Foundation of China ([2018]155).

### Author Contributions Statement

Conceived and designed the experiments: Yanbing Li & Zhi Li. Performed the experiments: Zhenyu Jiao & Yingqun Chen. Analyzed the data: Yang Xie. Contributed reagents/materials/analysis tools: Yang Xie. Wrote the paper: Zhenyu Jiao & Zhi Li. All authors reviewed the manuscript.

### Conflicts of Interest

The authors declare no conflict of interest. All authors read and approved the final manuscript.

